# A Novel Quantitative Metric Based on a Complete and Unique Characterization of Neural Network Activity: 4D Shannon’s Entropy

**DOI:** 10.1101/2023.09.15.557974

**Authors:** Sarita S. Deshpande, Wim van Drongelen

## Abstract

The human brain comprises an intricate web of connections that generate complex neural networks capable of storing and processing information. This information depends on multiple factors, including underlying network structure, connectivity, and interactions; and thus, methods to characterize neural networks typically aim to unravel and interpret a combination of these factors. Here, we present four-dimensional (4D) Shannon’s entropy, a novel quantitative metric of network activity based on the Triple Correlation Uniqueness (TCU) theorem. Triple correlation, which provides a complete and unique characterization of the network, relates three nodes separated by up to four spatiotemporal lags. Here, we evaluate the 4D entropy from the spatiotemporal lag probability distribution function (PDF) of the network activity’s triple correlation. Given a spike raster, we compute triple correlation by iterating over time and space. Summing the contributions to the triple correlation over each of the spatial and temporal lag combinations generates a unique 4D spatiotemporal lag distribution, from which we estimate a PDF and compute Shannon’s entropy. To outline our approach, we first compute 4D Shannon’s entropy from feedforward motif-class patterns in a simulated spike raster. We then apply this methodology to spiking activity recorded from rat cortical cultures to compare our results to previously published results of pairwise (2D) correlated spectral entropy over time. We find that while first- and second-order metrics of activity (spike rate and cross-correlation) show agreement with previously published results, our 4D entropy computation (which also includes third-order interactions) reveals a greater depth of underlying network organization compared to published pairwise entropy. Ultimately, because our approach is based on the TCU, we propose that 4D Shannon’s entropy is a more complete tool for neural network characterization.

**Author Summary:** Here, we present a novel entropy metric for neural network characterization, 4D Shannon’s entropy, based on triple correlation, which measures interactions among up to three neurons in time and space. Per the Triple Correlation Uniqueness (TCU) theorem, our 4D entropy approach is based on a complete and unique characterization of network activity. We first outline the method to obtain 4D Shannon’s entropy using a simulated spike raster of feedforward three-neuron configurations. We then apply this metric to an open-source, experimental dataset of rat cortical cultures over time to show that while first- and second-order interactions (spike rate and cross-correlation) show similar trends to published results, the TCU-based 4D Shannon’s entropy metric provides greater insights into later-stage network activity compared to the published pairwise entropy. As this metric is computed from a 4D distribution unique to the network, we propose that utilization of 4D entropy offers a clear advantage compared to currently utilized pairwise entropy metrics for neural network analyses. For this reason, neuroscientific and clinical applications abound – these may include analysis of distinct dynamical states, characterizing responses to medication, and identification of pathological brain networks, such as seizures.

## Introduction

Unraveling the intricate web of connections within neural networks remains a formidable task. As such, network characterization is an increasingly important frontier in neuroscience, continuously evolving to further uncover information hidden within the brain. Currently, there are multiple methods to characterize neural networks, and they typically do so by analyzing network dynamics and structure. Some of the commonly implemented methods include functional connectivity analysis, information theory, and network topology & graph theory analysis. Functional connectivity analyses involve untangling the temporal correlations and statistical dependencies between populations of neurons or brain regions [1, 2]. These methods investigate interactions among brain regions to decipher information flow and processing, identify regions of coordinated activity, and assess the strength of underlying network architecture [1, 2]. Researchers have investigated functional connectivity under different states, such as resting-state (evaluating spontaneous neural activity in the absence of external input) [3] and task- or state-based cases (assessing how network behaviors change in response to specific tasks or stimulation) [4].

Information theory methods quantify the amount of information contained within the network and typically employ metrics such as entropy and mutual information to do so [5, 6]. The goal of information theory is to assess how information is encoded and decoded [7]. Shannon’s entropy, as described in his consequential paper “A Mathematical Theory of Communication,” is a metric of the amount of uncertainty contained in a probability distribution function (PDF) [5]. Shannon’s entropy is defined as:

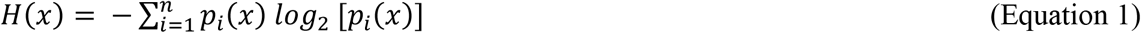

where p_i_ is the probability associated with state i, in our examples using spike raters x is a discrete variable, and log_2_ is the base-2 logarithm [5]. This measurement has since become a crucial element of information theory with applications to multiple fields, including computer science, economics, and biology. In addition, Shannon’s entropy has been used as a powerful tool in neuroscience, with respect to investigating neural information coding [8], functional connectivity [9, 10], complexity [11], neural variability [12], and information flow [13, 14].

Network topology and graph theory approaches aim to characterize networks based on their underlying structure and employ metrics such as clustering coefficient and average path length to provide insights into organization of network connectivity [15, 16]. One notable property used for these investigations is small-world networks [17], which are characterized by both high clustering and short path lengths (indicating rapid communication pathways which allow for efficient information processing) [17]. Small-world characteristics have also been used to investigate anatomical and functional connectivity of the brain [18, 19]. In addition to clustering coefficient and average path length, other network properties utilized in graph theory approaches include degree centrality and rich-club organization. Degree centrality measures the number of connections for a node within the network; a high degree centrality indicates a node with multiple connections, which may act as a “hub” or a crucial brain region for information processing [9, 20]. Building from this, rich club organization indicates that brain regions with high degree centrality form densely connected hubs, allowing for rapid information flow among brain regions [20, 21]. These metrics, as employed by network topology and graph theory approaches, analyze information processing in terms of underlying network architecture.

Adding to these network characterization approaches, we recently showed that third-order network motifs are sufficient to completely and uniquely characterize neural network activity [22]. To do so, we utilize triple correlation, which measures relationships among up to three nodes separated by four lags in time and space [22]. The purpose of this study is to build from the triple correlation approach and compute the entropy of a network from its spatiotemporal lag distribution. We apply this method to simulated spike rasters as well as a real-world experimental dataset. We show evidence that, since the triple correlation uniquely represents the underlying activity, the entropy based on triple correlation provides a more complete representation of the neuronal activity patterns and is therefore a compelling description of underlying network function. A short description of this work has appeared previously as an abstract in a conference proceeding [23].

## Methods

To add to the repertoire of network characterization and analysis methods, here we present 4D Shannon’s entropy. This novel metric is based on the Triple Correlation Uniqueness (TCU) theorem, which states that triple correlation provides a complete and unique characterization of the network [22]. In order words, there is a one-to-one relationship between the information contained in the network and the information contained in triple correlation [22]. Briefly, triple correlation is a signal processing tool that relates three nodes: one reference node and up to two other nodes separated by up to two lags in both space (n_1 &_ n_2_) and time (t_1 &_ t_2_). Given a spike raster (Fig 1A, [22]), r(n,t), where n is space and t is time, triple correlation (c_3_) is defined as:

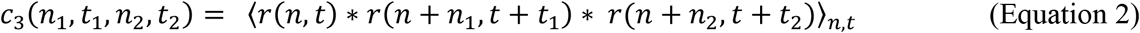

**Fig. 1.**
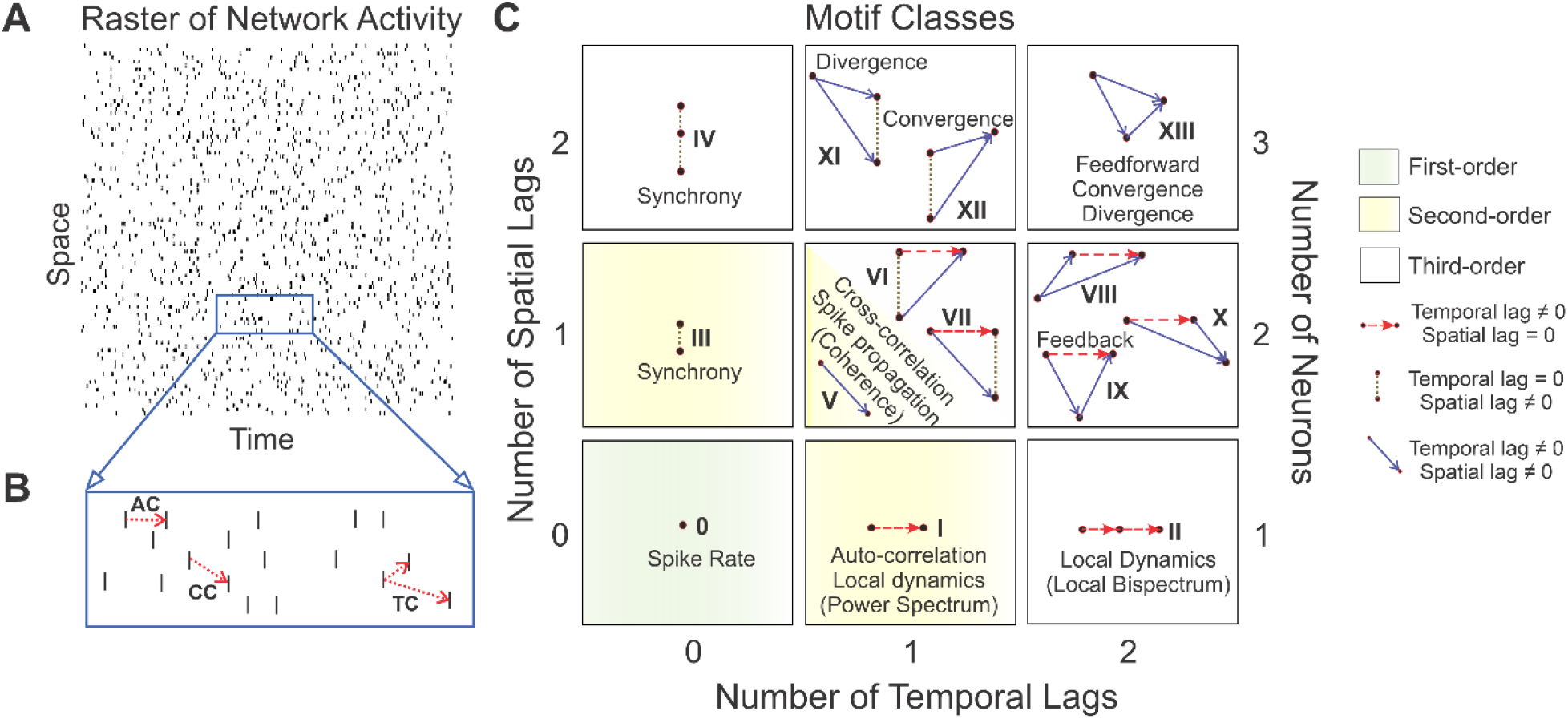
[22]: Triple correlation relates three nodes separated by up to two spatial and two temporal lags. **(A)** Given a spike raster, **(B)** autocorrelation (AC) relates two nodes separated by one temporal lag, cross-correlation (CC) relates two nodes separated by one spatial and one temporal lag, and triple correlation (TC) relates three nodes separated by up to two spatial and two temporal lags. These three-node configurations can be collapsed into **(C)** fourteen motif classes (0-XIII) which can embody well-studied neuronal processing properties. The fourteen motif classes are termed the motif-class spectrum. *Figure reprinted with permission from authors and under the terms of the Creative Commons CC BY license from [22]*.

In a binary spike raster consisting of 0s (no activity) and 1s (spikes), we can determine all non-zero contributions and their associated lags (n_1_, t_1_, n_2_, t_2_). This generates 169 possible three-node (motif) configurations, including those in which nodes overlap as in auto- and cross-correlation (Fig. 1B, [22]). These 169 motif configurations can be collapsed into fourteen qualitatively distinct motif classes based on symmetries that occur in time and space (Fig. 1C, [22]). These motif classes can embody well-studied neuronal processing properties, such as synchrony (motif classes III-IV), feedback (motif class IX), divergence (motif class XI), convergence (motif class XII), and feedforward (motif class XIII). We have previously developed a metric of quantifying network structure using triple correlation: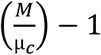, in which M is the observed prevalence of motif classes and μ_c_ is the conditioned theoretical expectation value which is takes into account the prevalence of first-order (motif class 0: spike rate; green highlighted box in Fig. 1C) and second-order motif classes (motif class I: autocorrelation; motif class III: synchrony; and motif class V: cross correlation; yellow highlighted areas in Fig. 1C) [22]. For example, computing μ_c_ for motif class VI is based on the prevalences of motif classes 0 (spike rate), I (autocorrelation), III (synchrony), and V (cross-correlation) (as motif class VI is comprised of each of these lower-order classes). Hence, values of 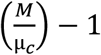 greater than 0 indicate the presence of network structure whereas values of 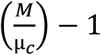 less than 0 indicate network activity comparable to chance levels [22]. Network activity across each of these fourteen motif classes is termed the motif-class spectrum.

Here, we build from triple correlation to compute the 4D entropy of a network from its unique 4D spatiotemporal lag distribution. Computing triple correlation across the network yields contributions at specific spatial and temporal lag combinations. If the contribution to the triple correlation is nonzero, we store both the contribution itself as well as the spatial and temporal lag combination that determined that contribution. After iterating over the entire network, we sum the contributions across each of the spatial and temporal lag combinations, generating a 4D spatiotemporal distribution for lags n_1_, t_1,_ n_2_, and t_2_. As per the TCU theorem [22], this 4D distribution is not only unique to its triple correlation (c_3_), but also unique to the network. In other words, the information contained in the 4D distribution is also contained in network and its triple correlation. From this distribution, we estimate a 4D probability distribution function (PDF). From these PDFs, Shannon’s entropy can be computed using Equation 1 (note that the entropy is 4D due to the four spatiotemporal lags).

## Results

To illustrate our approach, we determined the four-dimensional (4D) entropy of a network activity pattern of a simulated spike raster with a set of feedforward motif-class patterns (Fig. 2A; motif class XIII from Fig. 1). We computed the triple correlation by iterating through each spatial lag from −5:5 channels and each temporal lag from −4:4 bins (these lags were chosen arbitrarily for the sole purpose of outlining the method). We also determined the triple correlation for a set of spike-rate matched surrogate rasters, in which all spikes are shuffled over time and space (Fig. 2B, n=100 iterations). Summing the contributions yielded a 4D spatiotemporal lag distribution, of which a slice at n_2_ =0 & t_2_ =0 is shown in Fig. 2C-D. Note that within this slice at n_1_ =0 & t_1_ =0, the summed contribution to the triple correlation was 48, which is equivalent to the number of spikes in both the feedforward network raster and surrogate raster. For the three non-zero lag scenarios, with lags that can be positive and negative (hence the spatiotemporal symmetry in Fig. 2C), we find 48/3 = 16 for each case. The average distribution across 100 iterations of the surrogate raster is shown in Fig. 2D. From this, we estimated a 4D PDF (Fig. 2E-F) for each raster type (the mean PDF corresponding to 100 iterations of the surrogate raster is shown in Fig. 2F). From these PDFs (Fig. 2E: the raster PDF; Fig. 2F: mean surrogate PDF), we computed Shannon’s entropy (Fig. 2G). From Fig. 2G, the entropy of the feedforward network was lower, as expected, given the isolated motif-class network structure, and the entropy generated from the surrogate distribution (box-and-whisker plot, n=100 iterations) was higher than that of the raster, as expected given that the network structure had been abolished. This ideal simulated raster of isolated feedforward motif classes was used to outline and validate our methodology. We then sought to test this approach on real-world experimental dataset.

**Fig. 2:**
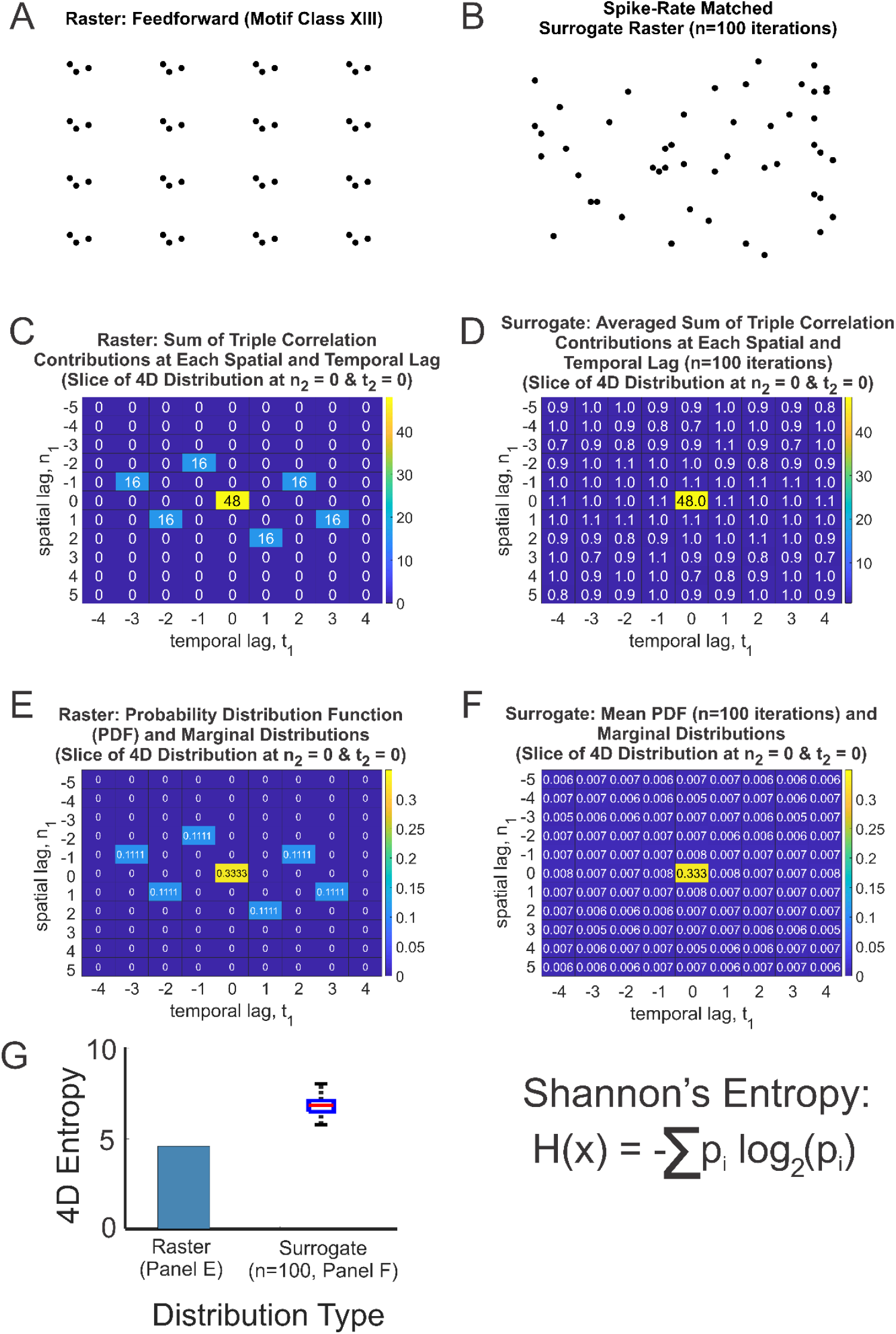
Overview of method to compute probability distribution function (PDF) and associated 4D entropy from the triple correlation of a simulated spike raster. **(A)** Given a spike raster of isolated feedforward motif class patterns and **(B**) one instance of a spike-rate matched surrogate (shuffled) raster (n=100 iterations), we compute triple correlation (using temporal lags from −4:4 bins and spatial lags from −5:5 bins for this example) and **(C-D)** sum the contributions to the triple correlation at each spatial and temporal lag combination for the feedforward network raster and the surrogate raster (average distribution for 100 iterations). **(E-F)** From this, we estimate the PDF. **(G)** 4D Shannon’s entropy computed for the raster and the whisker plot for the 100 surrogates.

We then applied our approach to an open-source dataset of spiking activity from microelectrode array recordings of rat cortical neurons [24, 25]. Briefly, cortical tissue was extracted from rat embryos (embryonic days 17-18) and plated on MEA well plates (12 wells per plate; 8×8 array of 64 electrodes per well) [24, 25]. Spiking activity was recorded from 2-35 days *in vitro* (DIV). Triple correlation was computed across 60 one-second epochs throughout the spike raster (n=12 wells) using spatial lags that cover all 64 electrodes and temporal lags from −50:50 ms. This temporal window of 100 ms covers monosynaptic excitatory (AMPA-& peak of NMDA-mediated) postsynaptic potentials and inhibitory (GABA_A_-mediated) postsynaptic potentials.

Comparing the published spike rate (Fig. 3A) with the spike rate computed by triple correlation (motif class 0, Fig. 3B) as well as the published pairwise (2D) correlated spectral entropy (CorSE) (Fig. 3C) with the spike-rate matched prevalence of motif class V (cross-correlation - a pairwise, 2D metric of spike propagation) (Fig. 3D) show similar trends in that the activity gradually increases, peaks around 24-28 DIV, and decreases from 31-35 DIV. The spatiotemporal lag PDF and its associated 4D entropy (Fig. 3E) were computed using triple correlation, which also yielded the fourteen individual motif-class profiles (Fig. 3F). The 4D entropy (Fig. 3E) shows no decrease from 31-35 DIV, as observed by the trends in the first-order and second-order metrics (Fig. 3A-D). This difference between the published 2D entropy and our 4D entropy can be explained by the trends in the third-order motifs. Motif classes VII-X initially increase as the other motif classes, but without a clear decrease around 31-35 DIV. Furthermore, motif classes II, XI, and XII follow the trend of increasing, peaking around 24-28 DIV, and then decreasing from 31-35 DIV, but not to 0 (Fig. 3F). The thick blue line in Fig. 3F indicates the median if the number of contributing wells > 7 (out of 12 total wells) (line not shown for DIV 2 and DIV 7 for motif classes VI and VII, due insufficient spike activity).

**Figure 3:**
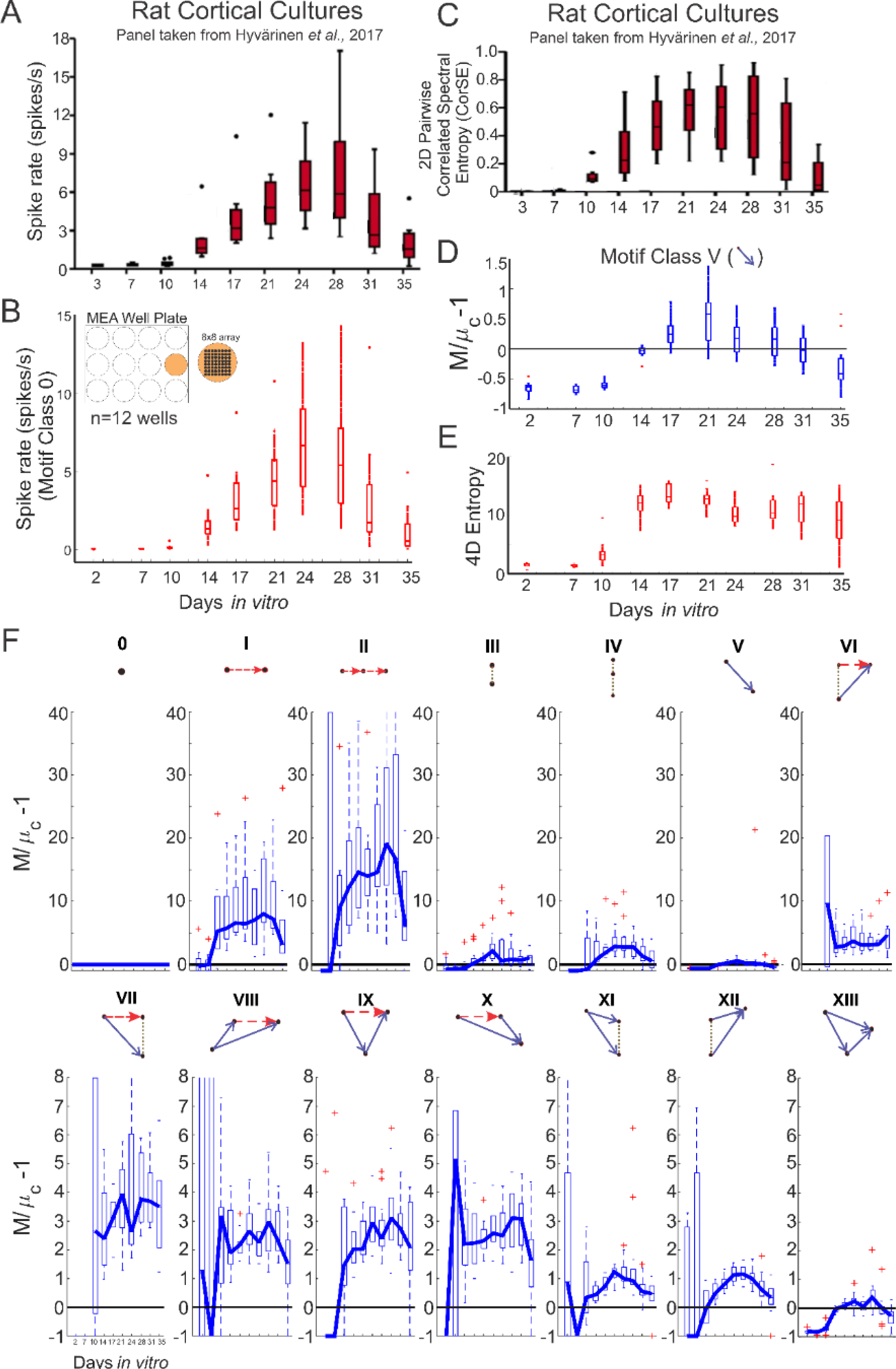
Characterizing network activity of rat cortical cultures over time using 4D entropy and the motif-class spectrum. **(A)** The spike rate from the published data **(B)** and triple correlation (motif class 0) show similar trends from 2-35 days *in vitro* (DIV). **(C)** The 2D correlated spectral entropy (CorSE) from the published data as well as **(D)** the prevalence of motif class V (pairwise 2D spike propagation) also show comparable trends. **(E)** The 4D entropy computation based on triple correlation shows a comparable trend from 2-17 DIV but does not decrease from 21-35 DIV, even though the spike rate **(A-B)** and second-order pairwise metrics **(C-D)** show this decrease from 31-35 DIV. **(F)** The fourteen motif-class profiles from triple correlation provide insight into individual network patterns. Motif classes V (cross-correlation; same as Panel D) and XIII (feedforward spike propagation) show similar trends to that of the published 2D CorSE values in that they gradually increase, peak around 24-28 DIV, and then decrease to network activity comparable to chance. Motif classes II, XI-XII also peak around 24-28 and decrease from 31-35 DIV, but do not revert to activity comparable to chance from 31-35. Boxes in the box-and-whisker plots represent the 25^th^, 50^th^ (median), and 75^th^ percentiles, and the whiskers indicate the range of the data.

## Discussion

Here, we present a novel method to compute the entropy of a neural network based on the 4D spatiotemporal probability distribution function (PDF) generated from triple correlation. This 4D entropy metric is based on the triple correlation spatiotemporal lag PDF, which captures network activity full and uniquely as leveraged by the recently described application of the Triple Correlation Uniqueness theorem to network analysis [22]. We first illustrate our approach on a spike raster with isolated feedforward motif-class patterns. We then apply this approach to an open-source dataset of spiking activity recorded from rat cortical cultures and demonstrate that the 4D entropy value captures underlying network activity from 31-35 DIV, whereas the published 2D entropy shows a decrease. In addition, the trends for motif classes 0 (spike rate) and V (cross-correlation) align with the previously reported spike rate and the pairwise 2D correlated spectral entropy values. These trends also agree with networks in hippocampal cell cultures, in which excitatory synaptic density peaks between 14-20 DIV in sparse cultures and between 8-14 DIV for dense cultures [26]. This example presents compelling evidence that the 4D approach can create a more complete representation of the network patterns than the 2D one. Therefore, we propose a dual presentation for neural network characterization: 1) 4D entropy as a metric of overall network activity and 2) the fourteen motif-class profiles as metrics of individual network patterns underlying structure.

Like most signal processing tools, there are limitations to the application and interpretation of the 4D entropy metric presented here. The sampling of the network as well as the noise embedded within the data are two important criteria to consider prior to analysis: does the data sufficiently represent the network, and is the signal obscured by noise? These questions, however, are not unique to computing triple correlation or 4D entropy – they should be accounted for with any analysis pipeline.

As previously described, TCU-based approaches (including the novel 4D entropy metric) have broad applicability in network analysis, both within and outside neuroscience [22]. For neuronal network analyses, this is valid for analysis of the ongoing dynamics reflected in local field potentials, EEG, calcium imaging, and fMRI. One critical adaptation required in these cases is to account for theoretical expectations of randomized controls. For instance in spike rasters, the statistics of point processes provides one mechanism for control (as outlined in [22]), whereas in EEG one might, for example, use statistics of colored noise instead. An additional aspect to consider is the spatial organization of the network. For example, in our examples (Figs. 2 and 3), the spatial ordering of the neurons in the spike raster is arbitrary in the sense that we assume that each channel may interact with every other channel within the spatial lag limitations. However, this assumption may not withstand for other data types, such as in EEG, where the spatial map of electrodes will affect ordering and spatial lags to be considered.

Although it is essential to organize triple correlation-based analyses according to data type and hypotheses, future applications of its applications abound. The TCU and the 4D entropy metric can be used to identify distinct dynamical states as well as to map neural networks at multiple scales. Furthermore, the 4D distribution (which is unique to the network as per the TCU) provides a rich source for network analysis beyond just the motif-class spectrum and 4D entropy. For instance, further insight into a network’s function can be obtained from analyzing the 4D distribution to identify regions or hubs of interest based on the predominant temporal lag combinations, which may point to specific connectivity and neurotransmitter systems involved, or the predominant spatial lag pointing to the regions associated with ongoing information processing. In addition, the Fourier transform of the triple correlation is the bispectrum [22]. As such, generating a 4D distribution based on the frequency components (unique to the bispectrum as per the TCU) can also indicate important relationships across oscillatory components of different frequencies. For example, in EEG, this may include analysis of nonlinear interactions among frequency components, and in the case of epilepsy, classification of epileptic states [27] and prediction of epileptic seizures [28, 29].

## Code Availability

The figures presented here were generated using Matlab 2023b (Mathworks) All scripts and functions used for analysis and generation of figures are available on the following Github repository: https://github.com/ssdeshpande334/4D_Entropy_Triple_Correlation. The corresponding triple correlation & 4D entropy output files are available on: https://zenodo.org/record/8347479 (DOI: 10.5281/zenodo.8347479).

## Data Availability

The open-source experimental dataset for Figure 3 is available from [25].

## Acknowledgements

S.S.D. is supported by F31NS127493. W.v.D is supported by R01 NS084142-02.

